# African-lineage Zika virus replication dynamics and maternal-fetal interface infection in pregnant rhesus macaques

**DOI:** 10.1101/2020.11.30.405670

**Authors:** Chelsea M. Crooks, Andrea M. Weiler, Sierra L. Rybarczyk, Mason Bliss, Anna S. Jaeger, Megan E. Murphy, Heather A. Simmons, Andres Mejia, Michael K. Fritsch, Jennifer M. Hayes, Jens C. Eickhoff, Ann M. Mitzey, Elaina Razo, Katarina M. Braun, Elizabeth A. Brown, Keisuke Yamamoto, Phoenix M. Shepherd, Amber Possell, Kara Weaver, Kathleen M. Antony, Terry K. Morgan, Dawn M. Dudley, Eric Peterson, Nancy Schultz-Darken, David H. O’Connor, Emma L. Mohr, Thaddeus G. Golos, Matthew T. Aliota, Thomas C. Friedrich

## Abstract

Following the Zika virus (ZIKV) outbreak in the Americas, ZIKV was causally associated with microcephaly and a range of neurological and developmental symptoms, termed congenital Zika syndrome (CZS). The isolates responsible for this outbreak belonged to the Asian lineage of ZIKV. However, in-vitro and in-vivo studies assessing the pathogenesis of African-lineage ZIKV demonstrated that African-lineage isolates often replicated to high titer and caused more severe pathology than Asian-lineage isolates. To date, the pathogenesis of African-lineage ZIKV in a translational model, particularly during pregnancy, has not been rigorously characterized. Here we infected four pregnant rhesus macaques with a low-passage strain of African-lineage ZIKV and compared its pathogenesis to a cohort of four pregnant rhesus macaques infected with an Asian-lineage isolate and a cohort of mock-infected controls. Viral replication kinetics were not significantly different between the two experimental groups and both groups developed robust neutralizing antibody titers above levels considered to be protective. There was no evidence of significant fetal head growth restriction or gross fetal harm at delivery in either group. However, a significantly higher burden of ZIKV vRNA was found in maternal-fetal interface tissues in the macaques exposed to an African-lineage isolate. Our findings suggest that ZIKV isolates of any genetic lineage pose a threat to women and their infants.

**IMPORTANCE:** ZIKV was first identified over 70 years ago in Africa, but most of our knowledge of ZIKV is based on studies of the distinct Asian genetic lineage, which caused the outbreak in the Americas in 2015-16. In its most recent update, the WHO stated that improved understanding of African-lineage pathogenesis during pregnancy must be a priority. Recent detection of African-lineage isolates in Brazil underscores the need to understand the impact of these viruses. Here we provide the first comprehensive assessment of African-lineage ZIKV infection during pregnancy in a translational non-human primate model. We show African-lineage isolates replicate with similar kinetics to Asian-lineage isolates and are capable of infecting the placenta. However, there was no evidence of more severe outcomes with African-lineage isolates. Our results highlight both the threat that African-lineage ZIKV poses to women and their infants and the need for future epidemiological and translational in-vivo studies with African-lineage ZIKV.

## INTRODUCTION

Zika virus (ZIKV) gained global notoriety in 2015 when it caused a large epidemic of febrile illness in the Americas and, for the first time, was causally associated with birth defects in infants born to mothers who became infected while pregnant (1). Why was ZIKV, which was first isolated in Uganda in 1947, not causally linked to birth defects prior to this outbreak in the Americas? Several hypotheses have emerged to explain why congenital ZIKV infection seems like a new complication, including that women in Africa are exposed to and therefore immune to the virus before childbearing age; that ZIKV circulating in the Americas acquired mutations that increased its ability to cause congenital ZIKV syndrome (CZS); or that ZIKV disease is enhanced by prior flavivirus immunity.

ZIKV circulates as two genetic lineages: African and Asian. The vast majority of animal model and epidemiological studies of ZIKV to date have focused on Asian-lineage viruses because they were responsible for the outbreak in the Americas. Therefore, relatively little is known about the pathogenic potential of African-linage viruses, particularly with regard to fetal outcomes. In cell culture experiments, African-lineage ZIKV isolates have been shown to replicate to higher titers and induce increased cell lysis as compared to Asian-lineage ZIKV (2–6). Particularly notable was the demonstration that African-lineage ZIKV isolates cause more rapid and more severe cytopathic effects (CPE) than Asian-lineage isolates in human embryonic stem-cell derived trophoblasts, which are cells critical for the development of the placenta (3, 4). Furthermore, in both pregnant and non-pregnant immunocompromised mouse models. African-lineage isolates have consistently shown both increased mortality and increased fetal harm as compared to Asian-lineage strains (5–8). Several experiments have been conducted with African-lineage ZIKV isolates in non-pregnant macaques. Two of the isolates used in these studies have extensive passage histories in mice and therefore cannot be considered natural ZIKV isolates (9). Still, in one study the virus replicated in the rhesus macaque host, but not as robustly as Asian-lineage isolates. A second study showed no replication of the isolate in Mauritian cynomolgus macaques, while a third study showed replication comparable to Asian-lineage viruses and subsequent protection against heterologous challenge (10–12). A low-passage isolate has been used in several non-human primate models and showed modest replication when inoculated intrarectally and intravaginally, and robust replication when inoculated subcutaneously (13, 14).

In the July 2019 epidemiological update on ZIKV, the WHO identified the assessment of fetal outcomes following infection with African-lineage viruses as a priority (15). This is underscored by recent findings of African-lineage isolates in South America, including evidence of fetal harm in a non-human primate naturally exposed to an African strain of ZIKV most closely related to the prototype strain. MR766 (16, 17). While the ability of African-lineage ZIKV to infect the maternal-fetal interface and cause fetal harm has been rigorously studied in cell culture and immunocompromised mice, it remains unclear how translational these findings are to humans.

To address this gap, we aimed to assess the pathogenic potential of a low-passage African-lineage ZIKV isolate during pregnancy in our well-established non-human primate model of ZIKV (18, 19). Recently, we demonstrated that the low-passage and highly pathogenic African-lineage ZIKV strain ZIKV/Aedes africanus/SEN/DAK-AR-41524/1984 (ZIKV-DAK; BEI Resources. Manassas. VA) replicated to higher titer in maternal serum and caused significantly greater fetal harm as compared to Asian-lineage ZIKV in *Ifnar1*^-/-^ C57BL/6 mice (8). Notably, placental pathology in mice infected with ZIKV-DAK was more severe than in mice infected with an Asian-lineage virus. Since contemporary ZIKV isolates from Africa are not readily available, this strain is one of the most recent, low-passage isolates available for pathogenesis studies.

We infected four pregnant macaques with ZIKV/Aedes-africanus/SEN/DakAr41524/1984 (ZIKV-DAK) during the late first trimester, monitored fetal health and growth throughout pregnancy and assessed fetal outcomes (presence of vRNA, gross abnormalities) at delivery at gestational day 155, approximately 1.5 weeks prior to full term. We compare data from a cohort of four pregnant macaques infected with ZIKV-DAK to a cohort of four pregnant macaques infected with Zika-virus/H.sapiens-tc/PUR/2015/PRVABC59_v3c2 (ZIKV-PR), a low-passage Asian-lineage isolate. This virus, isolated from a human in Puerto Rico in 2015, has been well characterized in rhesus macaques (10, 18–22). Although we did not find evidence of more severe fetal outcomes following infection with an African-lineage virus as compared to Asian-lineage virus, data from this study supports the hypothesis that ZIKV of both African- and Asian-lineage pose a threat to women and their infants.

## RESULTS

### ZIKV-DAK replicates to high titer in macaques and with similar replication to ZIKV-PR

Four pregnant rhesus macaques (*Macaca mulatta*) were subcutaneously inoculated with 10^4^ PFU of ZIKV-DAK between gestation day 45 and 50, late in the first trimester (Figure 1). The first trimester is associated with the greatest risk of CZS in pregnant women and is both a time of active neurological development and a time before which many women know that they are pregnant (23, 24). Following inoculation, plasma was collected daily for 10 days post infection (dpi), then twice weekly until viremia resolved, and then once weekly for the remainder of gestation. Negative viremia was defined as two consecutive timepoints with viral loads below the limit of quantification of our ZIKV QRT-PCR assay (100 copies/mL plasma). Virus replicated to high levels (10^5^-10^6^ vRNA copies/mL) in all four macaques, with viremia persisting through day 10 for all four macaques; two macaques had prolonged viremia out to 28-77 dpi (Figure 2A). When compared to a cohort of four macaques infected with ZIKV-PR using the same inoculation and sampling regimen, there were no statistically significant differences in viremia peak, duration, or area under the curve, suggesting that this African-lineage isolate replicates in pregnant macaques with similar kinetics to Asian-lineage isolates (Figure 2B).

**Figure 1.**
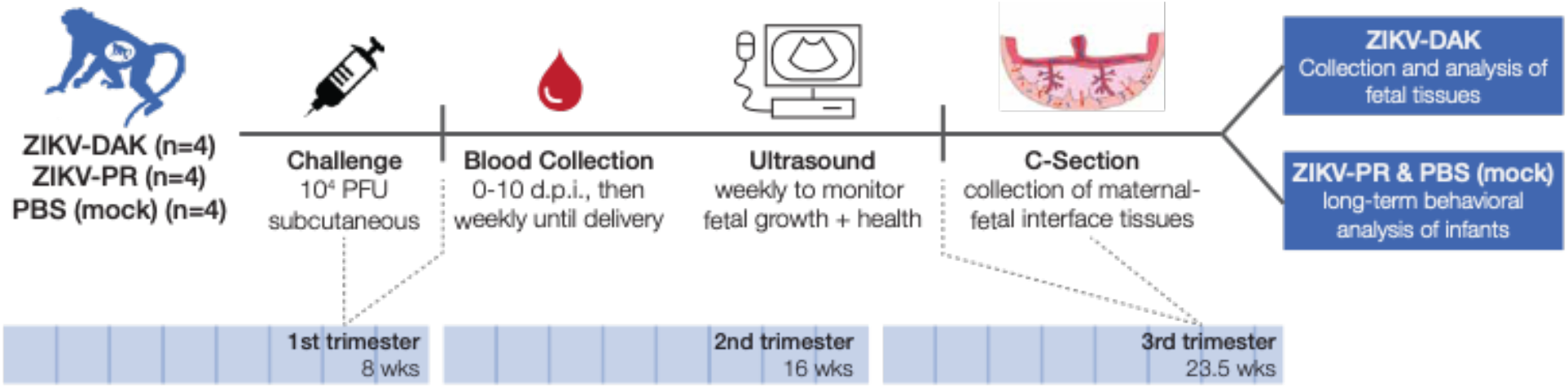
Study overview. Groups of four pregnant macaques were challenged between gestation days (gd) 45 and 50 (late first trimester) with either ZIKV-DAK, ZIKV-PR, or PBS (mock). Following viral challenge, blood was collected daily from 0-10 d.p.i., then twice weekly until viremia resolved, and then once weekly until delivery. Ultrasounds were performed once weekly to measure fetal health and growth. Between gd 155-160 (1.5 weeks prior to full term), infants were delivered via cesarean section, and maternal-fetal interface tissues, including the placenta, fetal membranes, umbilical cord, and placental bed were collected. Infants born to dams inoculated with ZIKV-DAK were humanely euthanized, and a comprehensive set of tissues were collected. Infants born to dams challenged with ZIKV-PR or PBS (mock) were paired with their mothers and followed for long-term behavioral analysis.

**Figure 2.**
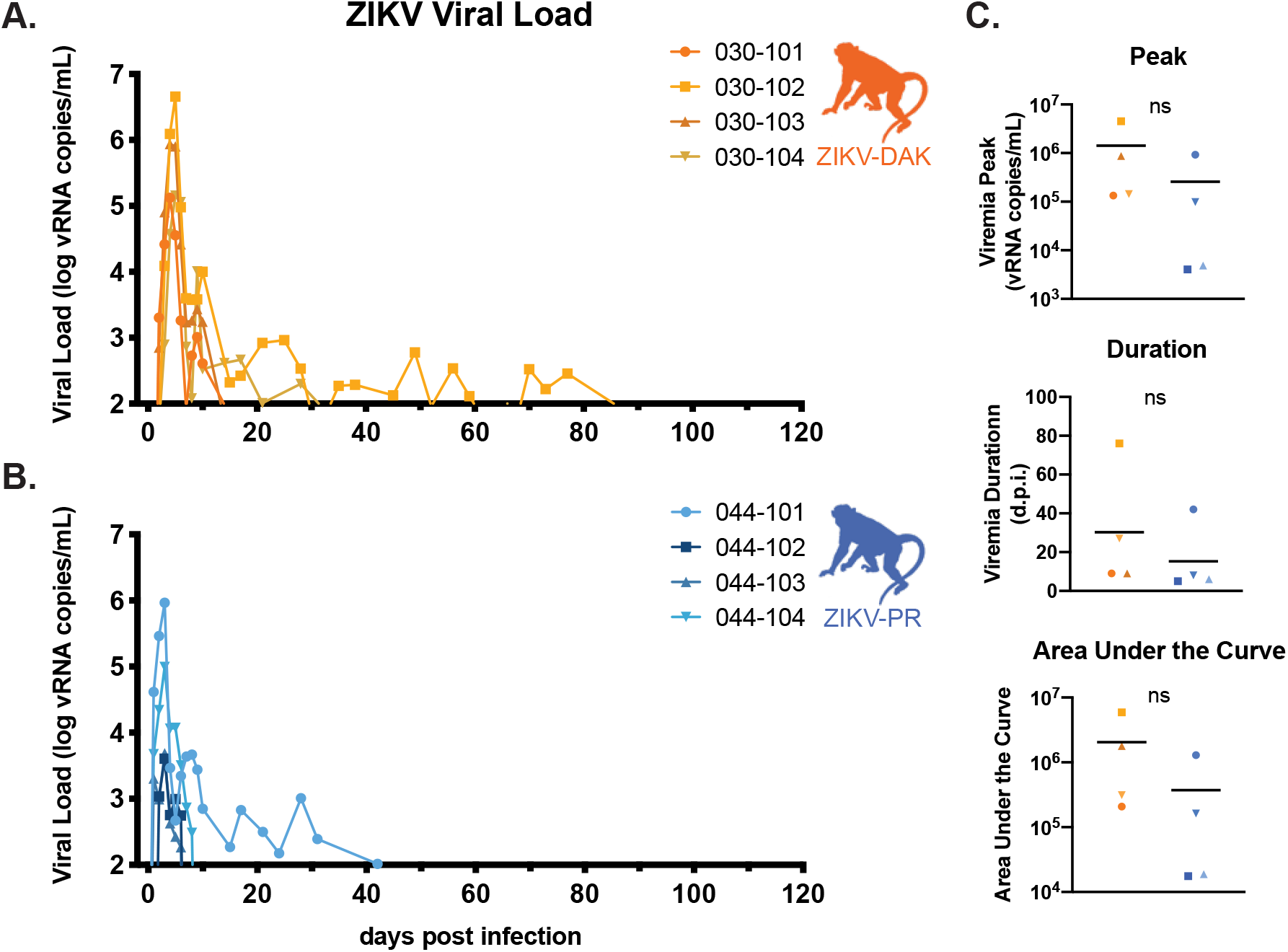
ZIKV-DAK and ZIKV-PR Viral replication kinetics. **A-B.** Viral load was determined using ZIKV-specific QRT-PCR of RNA isolated from plasma. Only values above the assay’s limit of quantification (100 copies/ml) are shown. **C.** There were no statistically significant differences in the peak, duration, or area under the curve of viremia between the two groups (two-sample t tests).

### ZIKV-DAK induces a robust nAb response

By 28 dpi, all macaques infected with either ZIKV-DAK or ZIKV-PR, regardless of viremia duration, had developed a robust neutralizing antibody titer (Figure 3A). The PRNT50 and 90 titers developed in response to ZIKV-DAK infection are significantly higher than those developed in response to ZIKV-PR infection (Figure 3B). The PRNT90 titers in macaques exposed to ZIKV-DAK are greater than the titers of macaques in a different study infected with the mouse-adapted isolate ZIKV-MR766, which were shown to be protective against heterologous challenge (10). Therefore, we expect that the immune response produced in these macaques infected with ZIKV-DAK during pregnancy would be protective against secondary ZIKV challenge.

**Figure 3.**
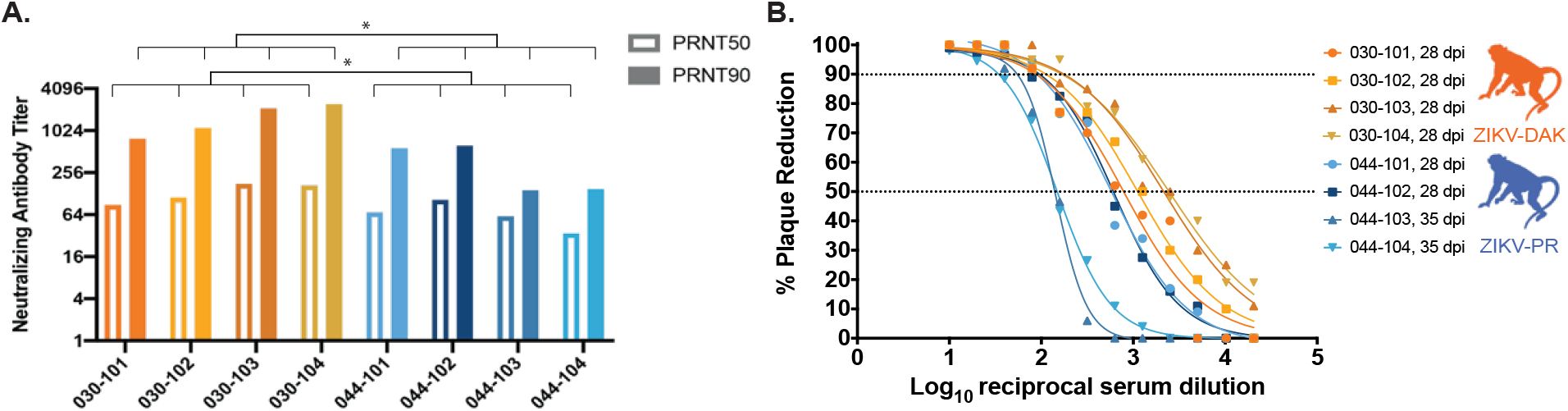
Neutralizing antibody titers. Plaque reduction neutralization tests (PRNT) were performed on serum samples collected between 28-35 days post infection to determine the titer of ZIKV-specific neutralizing antibodies. Neutralization curves were constructed using GraphPad Prism. PRNT90 and PRNT50 values were estimated using non-linear regression analysis and are shown on the bar graph in (**A**) and indicated with dotted lines in (**B**). PRNT50 and 90 titers were compared using an unpaired, parametric t-test. ZIKV-DAK-infected macaques had significantly higher PRNT50 (p=0.0371) and PRNT90 (p=0.0243) titers than ZIVK-PR-infected macaques.

### No reduction in fetal growth during gestation

Sonographic imaging was conducted weekly beginning one week prior to inoculation to evaluate fetal health (heart rate), overall fetal growth (abdominal circumference, femur length) and head growth (head circumference, biparietal diameter). No gross fetal or placental anomalies were observed. Varying amounts of placental calcification were noted on ultrasound in all four macaques exposed to ZIKV-DAK; however, calcifications were also observed in macaques infected with ZIKV-PR and in mock-inoculated macaques, suggesting that these qualitative observations are normal for gestational age or unrelated to viral infection.

Femur length, abdominal circumference, head circumference, and biparietal diameter measurements were compared to normative data developed from n=85 cynomolgus and rhesus macaques at the California National Primate Research Center (25, 26). We calculated the number of standard deviations that each fetal measurement differed from the normative data (z-score) at that gestational age. A linear mixed effects model with animal-specific random effects was used to evaluate the change in the outcome measures between gestation days 50-160. Growth was quantified by calculating the slope parameters for each experimental group. We then compared fetal growth in each group both to the normative data and to each of the other groups (Figure 4). When compared to the normative data, mock-infected animals had significantly reduced biparietal diameter growth (p=0.0207), while ZIKV-PR and ZIKV-DAK had a very modest, but statistically significant increase in head circumference growth (p=0.0230; p=0.0179). All other values were not significantly different from the normative data. When each of the experimental groups were compared to the mock-infected group, there was no significant reduction or increase in any of the growth measurements in the experimental group, suggesting that infection with either lineage of ZIKV did not restrict or enhance fetal growth.

**Figure 4.**
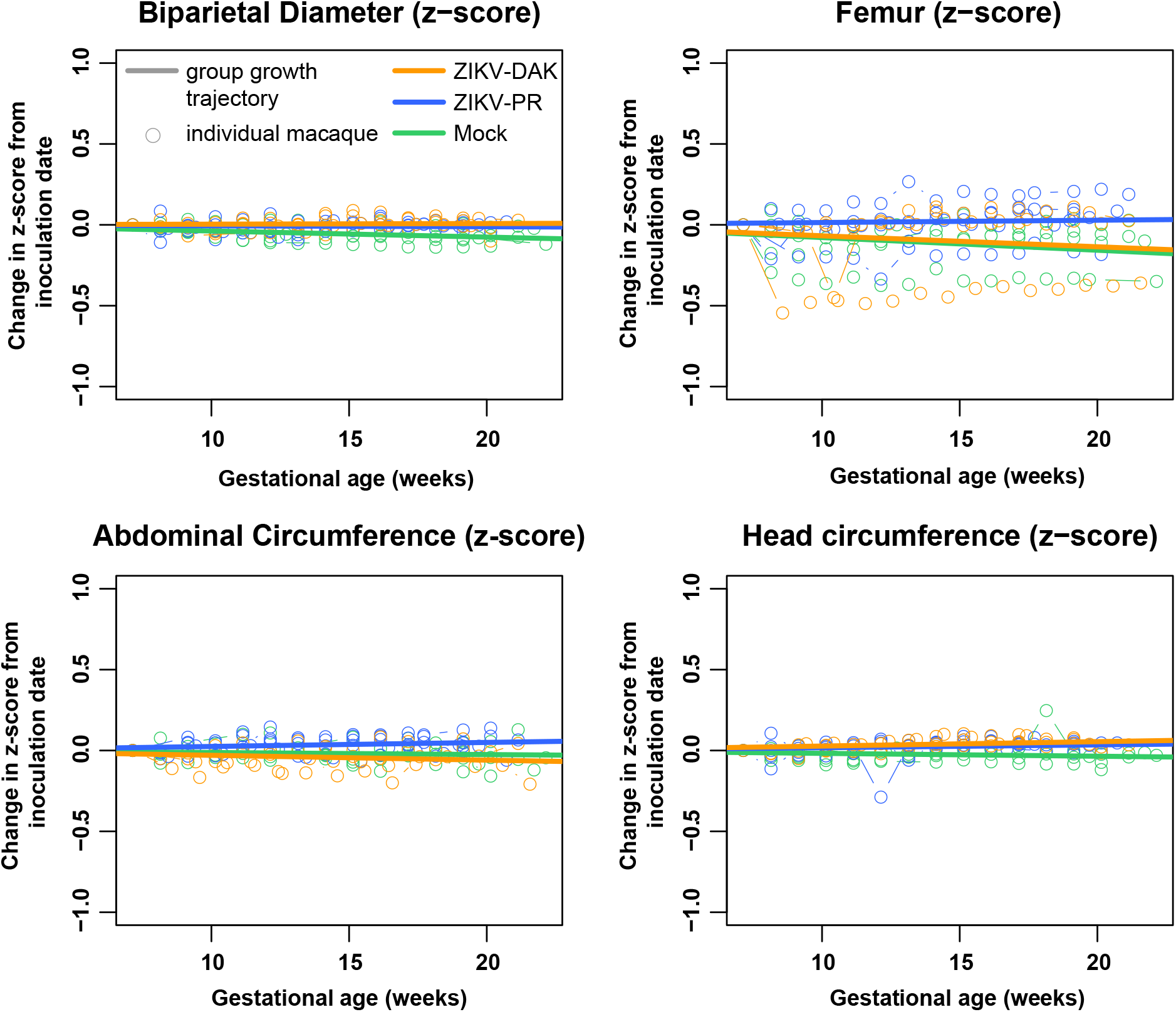
Intrauterine fetal growth. Sonographic imaging was performed weekly to measure fetal health and growth. Normative measurement data from the California National Primate Research Center was used to calculate z-scores for each weekly measurement for each macaque. The change in the z-score from the baseline measurement is plotted for each macaque with an open circle. Growth trajectories were quantified by calculating the regression slope parameters from baseline for each experimental group (solid line). When compared to the normative data, mock-infected animals had significantly reduced biparietal diameter growth (p=0.0207); ZIKV-PR and ZIKV-DAK had a very modest, but statistically significant increase in head circumference growth (p=0.0230; p=0.0179).

### No evidence of vertical transmission at delivery

At approximately gestation day 155 (full term is 165±10 days in rhesus macaques), fetuses of ZIKV-DAK-infected dams were delivered via cesarean section and humanely euthanized. No gross abnormalities were noted in any of the infants at delivery. A comprehensive set of maternal biopsies, maternal-fetal interface tissues, and fetal tissues were collected for vRNA measurements and histopathological analysis. In the fetus, emphasis was placed on collecting tissues that may be involved in transmission of the virus and tissues that are likely to be sites of ZIKV replication, including the central nervous system. Infants of ZIKV-PR-infected dams were delivered via cesarean section at approximately gestation day 160 and are being assessed for long-term neurodevelopmental sequelae. As a result, no fetal tissues were collected for comparison. No ZIKV RNA was detected in any of the fetal tissues collected at the time of delivery from the four ZIKV-DAK pregnancies (Supplemental table 2). Because pregnancies were allowed to go to term, we cannot exclude the possibility that ZIKV-DAK was vertically transmitted earlier in gestation but cleared from the fetus before delivery. Histopathological examination of fetal tissues revealed evidence of minimal to mild neutrophilic lymphadenitis in 3 of 4 ZIKV-exposed animals. Because we also observed neutrophilic lymphadenitis in four mock-infected animals that underwent the same experimental regimen, this inflammation may be a feature of normal development or from experimental procedures rather than viral infection.

### ZIKV is present in a variety of maternal-fetal interface tissues at delivery

Macaques typically have a bidiscoid placenta. To understand ZIKV distribution in the placenta, each placental disc was dissected into its individual cotyledons (perfusion domains), and a sample from the decidua, chorionic villi, and chorionic plate were taken from each cotyledon for both viral loads and histology (27). To understand ZIKV distribution in the maternal-fetal interface, additional samples were taken from the fetal membranes, uterine placental bed, and umbilical cord for both viral loads and histology. To assess the presence of virus in the dam, a biopsy of the mesenteric lymph node, liver, and spleen was taken from dams exposed to ZIKV-DAK for viral loads and histology. Only mesenteric lymph node biopsies were collected from dams exposed to ZIKV-PR. In the ZIKV-DAK dams, 3 of 12 biopsies, representing 2 different tissue types from 2 different macaques, were positive (Supplementary Table 2). In the ZIKV-PR dams, 1 of 4 mesenteric lymph node biopsies were positive.

ZIKV RNA was present in the placenta and maternal-fetal interface in all four ZIKV-DAK infected animals to varying degrees regardless of the duration of viremia (Figure 5A). The highest burden was found in the decidua (basalis), chorionic plate, and chorionic villi and at lower levels in the fetal membranes. No vRNA was identified in the placental bed of the uterus or umbilical cord tissues (Figure 5A) or in the amniotic fluid or umbilical cord blood (Supplemental Table 2). In contrast, there were fewer positive tissues in the maternal-fetal interface tissues of macaques infected with ZIKV-PR. All but one of the tissues positive for ZIKV-PR RNA were from a single macaque, 044-101. No vRNA was detected in the umbilical cord (Figure 5A), amniotic fluid, or umbilical cord plasma (Supplemental Table 2) in ZIKV-PR infected animals. When compared to the cohort of macaques infected with ZIKV-PR, there is a significantly greater burden of ZIKV vRNA present in the ZIKV-DAK cohort in the decidua, chorionic plate, and chorionic villi (Figure 5A). There were no significant differences in the vRNA burden in the fetal membranes, uterine placental bed, or umbilical cord.

**Figure 5.**
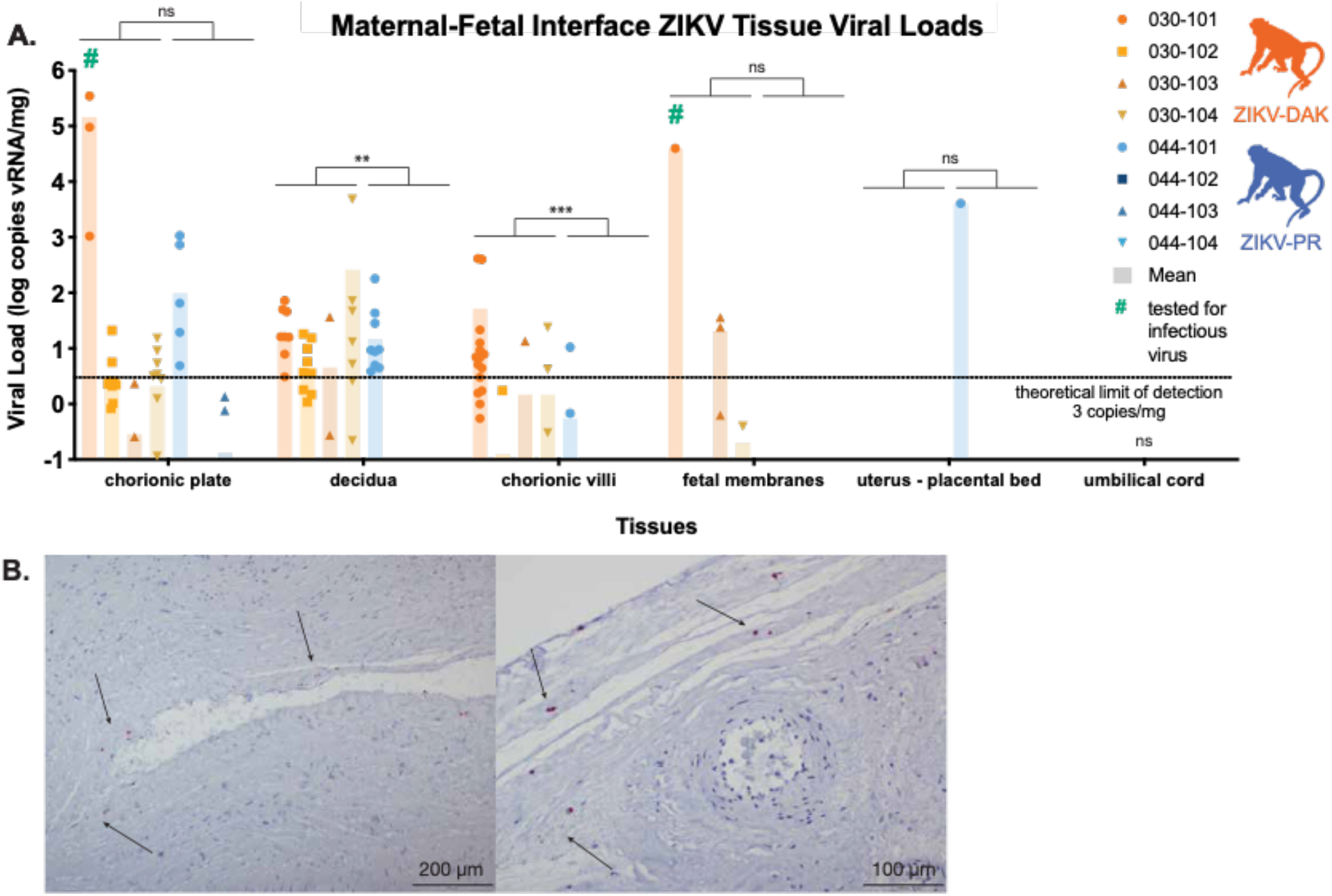
vRNA at the maternal-fetal interface. For each macaque, tissue biopsies were collected from the chorionic plate, chorionic villi, and decidua from each placental cotyledon; one to three biopsies were collected from the fetal membranes; and one biopsy was collected from the uterine placental bed and umbilical cord. Viral load was determined by ZIKV-specific QRT-PCR from RNA isolated from tissue samples. **A.** Viral load of maternal-fetal interface tissues. A non-parametric Mann-Whitney test was used to assess statistical significance between each experimental group in samples containing more than the theoretical limit of detection of 3 copies vRNA/mg tissue (**p<0.01, ***p<0.001). **B.** Representative images of in situ hybridization performed on fixed tissue sections from each of the placental cotyledons from 030-101. Positive staining for ZIKV RNA (red, arrows) was identified in 11 of the 17 cotyledons tested, primarily in the chorionic plate.

To assess whether there was replicating virus present in the placenta at delivery (105-113 days post infection), three high viral load (>10^3^ copies/mg) chorionic plate samples and one fetal membrane sample from 030-101 were tested for the presence of infectious virus using plaque assay. Three of the four tissues tested (two chorionic plates and one fetal membrane tissue) were positive for infectious virus via plaque assay (Supplemental Table 3). To further understand the distribution of vRNA within the placenta, tissue sections of the placental cotyledons from 030-101 were evaluated using in-situ hybridization (ISH). ISH probes for the ZIKV genome were used to identify ZIKV RNA in the tissue sections, 11 of the 17 cotyledons tested were positive for ZIKV RNA, primarily in the chorionic plate, which is consistent with both QRT-PCR and plaque assay results (Figure 5B).

### Comprehensive histological examination of placental tissues

To better understand the impact of in-utero ZIKV-DAK infection, maternal-fetal interface tissues were evaluated microscopically. Gross histopathological evaluation of the maternal-fetal interface tissues of ZIKV-DAK-exposed animals revealed a primary finding of transmural infarction of the central section of the placenta (Supplementary Figure 1). Transmural placental infarctions are areas of ischemic necrotic placental villi extending from the trophoblastic shell of the basal plate to the chorionic plate and are considered to be a result of a lack of oxygenated maternal blood flow, These infarctions were present in all four macaques infected with ZIKV-DAK. In contrast, this was observed in 2 of 4 macaques infected with ZIKV-PR and 1 of 4 mock-infected macaques.

Further histological analysis examined a cross-section of each of the individual placental cotyledons for the presence of chronic histiocytic intervillositis (CHIV), infarctions, villous stromal calcifications, and vasculopathy (Table 1). We also compared placental weights. There were no statistically significant differences in weight or pathological findings between the experimental and control groups for any of the features. The presence of infarctions, not often observed in mock-infected controls, could indicate that the pathology observed is a result of normal placental maturation and aging or a result of weekly anesthesia from experimental procedures. Regardless, these evaluations underscore the need to include mock-infected controls when evaluating tissues for viral pathogenesis. In order to quantitatively assess the pathologies present in the maternal-fetal interface, a central cross-section of each placental disc was scored for 22 functional features (Supplementary Table 4; Supplementary Figure 3). There were no statistically significant differences between either of the experimental groups and the mock-infected controls for any of the scored features; however, there was a trend toward increased chronic and acute villitis in the ZIKV-DAK exposed animals.

**Table 1.** Placental cotyledon pathology.

| Group | Dam | % CHIV+ cotyledons | Infarcted cotyledons/total cotyledons (%) | Villous stromal calcifications (present/absent) | Vasculopathy (present/absent) | Placental weight (g) |
| --- | --- | --- | --- | --- | --- | --- |
| Mock | 044-105 | 0.0 | 5.88 | Present | Absent | 111.08 |
|  | 044-106 | 0.0 | 12.5 | Present | Absent | 106.5 |
|  | 044-107 | 0.0 | 0.0 | Present | Present | 144.48 |
|  | 044-108 | 0.0 | 45.5 | Present | Absent | 122.92 |
| ZIKV-DAK | 030-101 | 0.0 | 43.8 | Absent | Absent | 131.54 |
|  | 030-102 | 0.0 | 0.0 | Absent | Absent | 111.4 |
|  | 030-103 | 0.0 | 66.7 | Absent | Present | 135.32 |
|  | 030-104 | 0.0 | 57.9 | Present | Present | 124.71 |
| ZIKV-PR | 044-101 | 0.0 | 25.0 | Present | Absent | 172.59 |
|  | 044-102 | 0.0 | 33.3 | Present | Absent | 123.87 |
|  | 044-103 | 0.0 | 0.0 | Absent | Absent | 134.49 |
|  | 044-104 | 0.0 | 18.2 | Absent | Absent | 120.48 |

## DISCUSSION

Here we provide the first comprehensive analysis of a low-passage, African-lineage ZIKV isolate in pregnant non-human primates. The data presented here demonstrate that this African-lineage ZIKV isolate is capable of robust replication in rhesus macaques. Infection induces a strong neutralizing antibody response, at or above titers that have been shown to be protective against subsequent challenge two years following primary challenge (28). Regular monitoring of fetal growth via ultrasound did not reveal any significant intrauterine growth restriction as compared to mock-infected animals. ZIKV infection of the placenta has been shown to be focal (21); therefore, in addition to assessment of well-established sequelae of viral infection at the maternal-fetal interface, we completed an extensive virological and histological evaluation of the placenta at delivery. Viral load testing of tissues from the extensive dissection of the placental discs into individual cotyledons and specific segments thereof revealed a higher burden of ZIKV in the chorionic plate in animals exposed to ZIKV-DAK, while plaque assay and ISH testing of high viral load samples confirmed the presence of infectious virus in the chorionic plate. ZIKV vRNA was also regularly found in the decidua and chorionic villi, and to a lesser extent in the fetal membranes. Despite a high burden of ZIKV in the chorionic plate – the fetal side of the placenta – there was no evidence of vertical transmission at delivery, even though infectious ZIKV was detected in the chorionic plate of one animal. This suggests that the vRNA burden in the maternal-fetal interface is not a robust predictor of clinical outcome for the fetus, but it does not preclude the possibility that infants may develop clinical sequelae later in life due to viral exposure, or placental insufficiency during gestation. Many normal-appearing infants exposed to ZIKV in utero develop neurodevelopmental delays in the years after birth (29–32).

This cohort of macaques infected with an African-lineage virus was compared to a cohort of macaques infected with an Asian-lineage virus and a mock-infected control group. Based on previous studies in cell culture and mice, we expected to see a more severe phenotype in the macaques that were infected with the African-lineage virus (2, 3, 3, 4, 4, 5, 5–8). We expected this more severe phenotype to manifest as enhanced viral replication (as determined by higher peak or longer duration of viremia), gross fetal abnormalities at delivery, or fetal demise. However, the only feature that was significantly different between the experimental groups was an increase in the burden of vRNA in the chorionic plate, chorionic villi and decidua.

To date, few studies of Asian-lineage viruses in non-human primates have shown clear evidence of fetal harm, despite a clear association between Asian-lineage ZIKV and CZS. A minority of human pregnancies known to be affected by ZIKV result in CZS (5-14%) or fetal loss (4-7%) (33); therefore, it is perhaps unsurprising that there is limited evidence of fetal harm in small non-human primate studies. In this study, ZIKV-DAK infection of pregnant macaques resembled infection of ZIKV-PR across several parameters, including infection of MFI tissues. Therefore, although we did not observe direct fetal harm, our findings suggest that African-lineage viruses have similar capacity to cause fetal harm as Asian-lineage viruses. This data suggests that African-lineage ZIKV poses a threat to women and their infants, which should be taken into account when providing public health guidance. While African-lineage ZIKV had been thought to be geographically confined to Africa, recent studies have identified African-lineage isolates in South America (16, 17). This highlights the need for continuing study of ZIKV of both genetic lineages.

A significant limitation of this study is the small sample size (n=4) in each of the experimental groups. Particularly when studying a pathogen whose most severe effects are only found in a minority of cases (33), modeling rare events in a small study is difficult and we cannot capture the full range of disease experienced by women infected with ZIKV during pregnancy. We also tested a single inoculation dose, virus strain, and inoculation time point; different experimental conditions may reveal different outcomes, which could include more stark differences between the lineages. Furthermore, this study focused on characterizing the pathogenesis of African-lineage ZIKV as compared to Asian-lineage but did not seek to understand the mechanisms of the vertical transmission of ZIKV or the potential mechanisms underlying differences between the lineages. Future studies should investigate these mechanisms and conduct more thorough epidemiological studies of African-lineage ZIKV, which may shed light on the reasons why ZIKV had not been associated with fetal harm prior to the outbreak in the Americas.

## METHODS

### Experimental design

This study was designed to assess the pathogenic potential of a low-passage African-lineage ZIKV isolate during pregnancy in a non-human primate model. Four pregnant Indian origin rhesus macaques (*Macaca mulatta*) were inoculated subcutaneously with 1×10^4^ PFU of ZIKV-DAK between 44-50 days of gestation (term is 165 ± 10 days). Macaques were monitored throughout the remainder of gestation. At approximately gestation day 155, infants were delivered via c-section and humanely euthanized. A comprehensive set of maternal biopsies, maternal-fetal interface and fetal tissues were collected for analysis. For the Asian-lineage group, four pregnant Indian origin rhesus macaques (*Macaca mulatta*) were inoculated subcutaneously with 1×10^4^ PFU of ZIKV-PR between 44-50 days of gestation (term is 165 ± 10 days). Macaques were monitored throughout the remainder of gestation. At approximately gestation day 160, infants were delivered via cesarean section and monitored for long-term development. A comprehensive set of maternal biopsies and maternal-fetal interface were collected for analysis. A cohort of four pregnant PBS-inoculated animals served as a control group and underwent the same experimental regimen, including the sedation for all blood draws and ultrasounds, as the ZIKV-PR cohort. Data used in this manuscript are publicly available at openresearch.labkey.com under study ZIKV-030 (ZIKV-DAK) and ZIKV-044 (ZIKV-PR and mock).

### Ethical approval

This study was approved by the University of Wisconsin College of Letters and Sciences and Vice Chancellor for Research and Graduate Education Centers Institutional Animal Care and Use Committee (Protocol numbers: G005401 and G006139).

### Care and use of macaques

All macaque monkeys used in this study were cared for by the staff at the WNPRC in accordance with the regulations and guidelines outlined in the Animal Welfare Act and the Guide for the Care and Use of Laboratory Animals and the recommendations of the Weatherall report (https://royalsociety.org/topics-policy/publications/2006/weatherall-report/). All macaques used in the study were free of *Macacine herpesvirus 1*, simian retrovirus type D (SRV), simian T-lymphotropic virus type 1 (STLV), and simian immunodeficiency virus. For all procedures (including physical examinations, virus inoculations, ultrasound examinations, and blood collection), animals were anaesthetized with an intramuscular dose of ketamine (10 mg/kg). Blood samples were obtained using a vacutainer system or needle and syringe from the femoral or saphenous vein.

### Cells and viruses

ZIKV/Aedes-africanus/SEN/DakAr41524/1984 (ZIKV-DAK) was originally isolated from *Aedes africanus* mosquitoes with a round of amplification on *Aedes pseudocutellaris* cells, followed by amplification on C6/36 cells and two rounds of amplification on Vero cells. ZIKV-DAK was obtained from BEI resources (Manassas, VA). Zika-virus/H.sapiens-tc/PUR/2015/PRVABC59_v3c2 (ZIKV-PR) was originally isolated from a human in Puerto Rico in 2015, with three rounds of amplification on Vero cells, was obtained from Brandy Russell (CDC, Fort Collins, CO, USA). African Green Monkey kidney cells (Vero; ATCC #CCL-81) were maintained in Dulbecco’s modified Eagle medium (DMEM) supplemented with 10% fetal bovine serum (FBS; Hyclone, Logan, UT), 2 mM L-glutamine, 1.5 g/L sodium bicarbonate, 100 U/ml penicillin, 100 μg/ml of streptomycin, and incubated at 37°C in 5% CO2. *Aedes albopictus* mosquito cells (C6/36; ATCC #CRL-1660) were maintained in DMEM supplemented with 10% fetal bovine serum (FBS; Hyclone, Logan, UT), 2mM L-glutamine, 1.5 g/L sodium bicarbonate, 100 U/ml penicillin, 100 μg/ml of streptomycin, and incubated at 28°C in 5% CO_2_. The cell lines were obtained from the American Type Culture Collection, were not further authenticated, and were not specifically tested for mycoplasma. Virus stocks were prepared by inoculation onto a confluent monolayer of C6/36 cells; a single, clarified stock was harvested for each virus, with a titer of 7.3 × 10^8^ PFU/ml for ZIKV-DAK and 1.58 × 10^7^ PFU/ml for ZIKV-PR. Deep sequencing with limited PCR cycles confirmed that the ZIVK-DAK virus stock was identical to the reported sequence in GenBank (KY348860) at the consensus level. Five nucleotide variants were detected at 5.1-13.1% frequency (Supplementary Table 1). PCR-free deep sequencing did not detect any evidence of Dezidougou virus, an insect-specific *Negevirus* that is present in some ZIKV-DAK stocks. Amplicon deep sequencing of ZIKV-PR virus stock using the methods described in Quick, et al. (34) revealed two consensus-level nucleotide substitutions in the stock as compared to the reported sequence in GenBank (KU501215), as well as seven other minor nucleotide variants detected at 5.3-30.6% frequency (Supplementary Table 1).

### Plaque Assay

All titrations for virus quantification from virus stocks and screens for infectious ZIKV from macaque tissue were completed by plaque assay on Vero cell cultures as previously described (35). Briefly, duplicate wells were infected with 0.1 ml aliquots from serial 10-fold dilutions in growth media and virus was adsorbed for one hour. Following incubation, the inoculum was removed, and monolayers were overlaid with 3ml containing a 1:1 mixture of 1.2% oxoid agar and 2X DMEM (Gibco. Carlsbad. CA) with 10% (vol/vol) FBS and 2% (vol/vol) penicillin/streptomycin (100 U/ml penicillin. 100 μg/ml of streptomycin). Cells were incubated at 37°C in 5% CO2 for four days for plaque development. Cell monolayers were then stained with 3 ml of overlay containing a 1:1 mixture of 1.2% oxoid agar and 2X DMEM with 2% (vol/vol) FBS. 2% (vol/vol) penicillin/streptomycin, and 0.33% neutral red (Gibco). Cells were incubated overnight at 37 °C and plaques were counted.

### Inoculations

Inocula were prepared from a viral stock propagated on a confluent monolayer of C6/36 cells. The stocks were thawed, diluted in PBS to 10^4^ PFU/ml and loaded into a 1 mL syringe that was kept on ice until challenge. Animals were anesthetized as described above and 1 ml of inocula was delivered subcutaneously over the cranial dorsum. Animals were monitored closely following inoculation for any signs of an adverse reaction.

### Ultrasound measurements

Ultrasound measurements were taken according to the procedures described previously (19). Briefly, dams were sedated with ketamine hydrochloride (10mg/kg) for weekly sonographic assessment to monitor the health of the fetus (heart rate) and to take fetal growth measurements, including the fetal femur length (FL), biparietal diameter (BPD), head circumference (HC), and abdominal circumference (AC). Weekly fetal measurements were plotted against mean measurement values and standard deviations for fetal macaques developed at the California National Primate Research Center (25, 26). Additional Doppler ultrasounds were taken as requested by veterinary staff.

Gestational age standardized growth parameters for fetal HC. BPD. AC, and FL were evaluated by calculating gestational age specific z-values from normative fetal growth parameters. Linear mixed effects modeling with animal-specific random effects was used to analyze the fetal growth trajectories with advancing gestational age. In order to account for differences in fetal growth parameters at the date of inoculation, changes in fetal growth parameters from date of inoculation (~day 50) were analyzed. That is, changes in fetal growth parameters from date of inoculation were regressed on gestational age (in weeks). An autoregressive correlation structure was used to account for correlations between repeated measurements of growth parameters over time. The growth trajectories were quantified by calculating the regression slope parameters which were reported along with the corresponding 95% confidence intervals (CI). Fetal growth was evaluated both within and between groups. All reported P-values are two-sided and P<0.05 was used to define statistical significance. Statistical analyses were conducted using SAS software (SAS Institute. Cary NC), version 9.4.

### Viral RNA isolation from blood

Viral RNA was isolated from macaque blood samples as previously described (18, 35). Briefly, plasma was isolated from EDTA-anticoagulated whole blood on the day of collection either using Ficoll density centrifugation for 30 minutes at 1860 x g if the blood was being processed for PBMC, or it was centrifuged in the blood tube at 1400 x g for 15 minutes. The plasma layer was removed and transferred to a sterile 15 ml conical and spun at 670 x g for an additional 8 minutes to remove any remaining cells. Viral RNA was extracted from a 300 μL plasma aliquot using the Viral Total Nucleic Acid Kit (Promega. Madison. WI) on a Maxwell 16 MDx or Maxwell RSC 48 instrument (Promega. Madison. WI).

### Viral RNA isolation from tissues

Tissue samples, cut to 0.5 cm thickness on at least one side, were stored in RNAlater at 4°C for 2-7 days. RNA was recovered from tissue samples using a modification of the method described by Hansen et al., 2013 (36). Briefly, up to 200 mg of tissue was disrupted in TRIzol (Lifetechnologies) with 2 x 5 mm stainless steel beads using the TissueLyser (Qiagen) for 3 minutes at 25 r/s twice. Following homogenization, samples in TRIzol were separated using Bromo-chloro-propane (Sigma). The aqueous phase was collected and glycogen was added as a carrier. The samples were washed in isopropanol and ethanol precipitated. RNA was fully re-suspended in 5 mM tris pH 8.0.

### Quantitative reverse transcription PCR (QRT-PCR)

vRNA isolated from both fluid and tissue samples was quantified by QRT-PCR as previously described(8). The RT-PCR was performed using the SuperScript III Platinum One-Step Quantitative RT-PCR system (Invitrogen, Carlsbad. CA) on a LightCycler 96 or LightCycler 480 instrument (Roche Diagnostics, Indianapolis, IN). Viral RNA concentration was determined by interpolation onto an internal standard curve composed of seven 10-fold serial dilutions of a synthetic ZIKV RNA fragment based on a ZIKV strain derived from French Polynesia that shares >99% similarity at the nucleotide level to the Puerto Rican strain used in the infections described in this manuscript.

### Statistical analysis of viral loads

Plasma viral load curves were generated using GraphPad Prism software. The area under the curve of 0-10 d.p.i, was calculated and a two-sample t-test was performed to assess differences in the peak, duration, and area under the curve of viremia between macaques infected with ZIKV-DAK and ZIKV-PR. To compare differences in the viral burden in the maternal-fetal interface, a non-parametric Mann-Whitney test was used to assess differences in each of the maternal-fetal interface tissues. GraphPad Prism 8 software was used for these analyses.

### Plaque reduction neutralization test (PRNT)

Macaque serum was isolated from whole blood on the same day it was collected using a serum separator tube (SST). The SST tube was centrifuged for at least 20 minutes at 1400 x g, the serum layer was removed and placed in a 15 ml conical and centrifuged for 8 minutes at 670 x g to remove any additional cells. Serum was screened for ZIKV neutralizing antibody utilizing a plaque reduction neutralization test (PRNT) on Vero cells as described in (37) against ZIKV-PR and ZIKV-DAK. Neutralization curves were generated using GraphPad Prism 8 software. The resulting data were analyzed by non-linear regression to estimate the dilution of serum required to inhibit 50% and 90% of infection.

### Cesarean section and tissue collection

Between 155-160 days gestation, infants were delivered via cesarean section and tissues were collected. The fetus, placenta, fetal membranes, umbilical cord, and amniotic fluid were collected at surgical uterotomy and maternal tissues were biopsied during laparotomy. These were survival surgeries for the dams. For fetuses born to dams infected with ZIKV-DAK, the fetus was euthanized with an overdose of sodium pentobarbitol (50 mg/kg) and the entire conceptus (fetus, placenta, fetal membranes, umbilical cord, and amniotic fluid) was collected and submitted for tissue collection and necropsy. For fetuses born to dams infected with ZIKV-PR, the infant was removed from the amniotic sac, the umbilical cord clamped, and neonatal resuscitation performed as needed. The placenta, amniotic fluid, and fetal membranes were then collected. Infants were placed with their mothers following the dam’s recovery from surgery.

Tissues were dissected as previously described(19) using sterile instruments that were changed between each organ and tissue type to minimize possible cross contamination. Each organ/tissue was evaluated grossly in situ, removed with sterile instruments, placed in a sterile culture dish, and sectioned for histology, viral burden assay, and/or banked for future assays. Sampling priority for small or limited fetal tissue volumes (e.g., thyroid gland, eyes) was vRNA followed by histopathology, so not all tissues were available for both analyses. A comprehensive listing of all specific tissues collected and analyzed is presented in Figure 5A (maternal-fetal interface tissues) and Supplementary Table 2 (maternal biopsies and fetal tissues). Biopsies of the placental bed (uterine placental attachment site containing deep decidua basalis and myometrium), maternal liver, spleen, and a mesenteric lymph node were collected aseptically during surgery into sterile petri dishes, weighed, and further processed for viral burden and when sufficient sample size was obtained, histology.

In order to more accurately capture the distribution of ZIKV in the placenta, each placental disc was separated, fetal membranes sharply dissected from the margin, weighed, measured, and placed in a sterile dish on ice. A 1-cm-wide cross section was taken from the center of each disc, including the umbilical cord insertion on the primary disc, and placed in 4% paraformaldehyde. Individual cotyledons, or perfusion domains, were dissected using a scalpel and placed into individual petri dishes. From each cotyledon, a thin center cut was taken using a razor blade and placed into a cassette in 4% paraformaldehyde. Once the center cut was collected, the decidua and the chorionic plate were removed from the remaining placenta. From each cotyledon, pieces of decidua, chorionic plate, and chorionic villi were collected into two different tubes with different media for vRNA isolation and for other virological assays.

### Histology

Following collection, tissues were handled as described previously (35). All tissues (except neural tissues) were fixed in 4% paraformaldehyde for 24 hours and transferred into 70% ethanol until processed and embedded in paraffin. Neural tissues were fixed in 10% neutral buffered formalin for 14 days until processed and embedded in paraffin. Paraffin sections (5 μm for all tissues other than the brain (sectioned at 8μm)) were stained with hematoxylin and eosin (H&E). Pathologists were blinded to vRNA findings when tissue sections were evaluated microscopically. Photomicrographs were obtained using a bright light microscope Olympus BX43 and Olympus BX46 (Olympus Inc., Center Valley, PA) with attached Olympus DP72 digital camera (Olympus Inc.) and Spot Flex 152 64 Mp camera (Spot Imaging) and captured using commercially available image-analysis software (cellSens DimensionR. Olympus Inc, and spot software 5.2).

### Placental Histology Scoring

Pathological evaluation of the cross-sections of each of the placental cotyledons were performed blinded to experimental condition. Each of the cross sections were evaluated for the presence of chronic histiocytic intervillositis (CHIV), infarctions, villous stromal calcifications, and vasculopathy. A three-way ANOVA was performed to assess statistical significance among groups for each parameter, including placental weight.

Two of three boarded pathologists, blinded to vRNA findings, independently reviewed the central cross section of each placental disc and quantitatively scored the placentas on 22 independent criteria. Six of the criteria are general criteria assessing placental function. 2 assess villitis, three criteria assessing the presence of fetal malperfusion, and 11 criteria assessing the presence of maternal malperfusion. The scoring system was developed by Dr. Michael Fritsch. Dr. Heather Simmons, and Dr. Andres Mejia. A summary table of the criteria scored and the scale used for each criterion can be found in Supplementary Table 3. Once initial scores were assigned, pathologists met to discuss and resolve any significant discrepancies in scoring. Scores were assigned to each placental disc for most parameters, unless the evaluation score corresponded to the function of the entire placenta.

For criteria that are measured on a quantitative scale, median scores and interquartile range were calculated for each experimental group. For criteria that were measured on a binary “present/not present” scale, the cumulative incidence in each experimental group was calculated as a frequency and a percentage. For quantitative criteria, a non-parametric Wilcoxon rank test was used to calculate statistical significance between each of the groups and between the mock-infected group and the two ZIKV-infected groups. For binary features. Fisher’s exact test was used to calculate statistical significance between each of the groups and between the mock-infected group and the two ZIKV-infected groups. To determine whether chronic villitis correlated with the criteria assessing fetal malperfusion and whether chronic deciduitis correlated with the criteria assessing maternal malperfusion, scores were adjusted to be on the same scale (i.e., converting measures on a 0-1 scale to a 0-2 scale) so that each parameter carried equal weight in the combined score. A non-parametric Spearman’s correlation was used to determine the correlation.

### In situ hybridization

In situ hybridization was conducted on cross sections of placental cotyledons as previously described(20). Briefly, tissues were fixed in 4% PFA, alcohol processed, and paraffin embedded. Commercial ISH probes against the ZIKV genome (Advanced Cell Diagnostics, Cat No. 468361, Newark, California, USA) were used. ISH was performed using RNAscope^®^ Red 2.5 Kit (Advanced Cell Diagnostics, Cat No. 322350) according to the manufacturer’s instructions.

## Data availability

All of the data used for figure generation and statistical analysis in this manuscript can be found at https://github.com/cmc0043/african-lineage-zikv-in-pregnant-macaques. In the future, primary data that support the findings of this study will also be available at the Zika Open Research Portal (https://openresearch.labkey.com/project/ZEST/begin.view). Data for the ZIKV-DAK infected cohort can be found under study ZIKV-030; data for ZIKV-PR and mock-infected cohorts can be found under ZIKV-044. Raw FASTQ reads (BioProject: PRJNA673500) and a FASTA consensus sequence (BioProject: PRJNA476611) of the challenge stock of ZIKV/Aedes africanus/SEN/DAK-AR-41524/1984 are available at the Sequence Read Archive. Raw FASTQ reads of the challenge stock of ZIKV PRVABC59 are available at the Sequence Read Archive, BioProject accession number PRJNA392686. The authors declare that all other data supporting the findings of this study are available within the article and its supplementary information files.

## ACKNOWLEDGEMENTS

We thank the Veterinary Services, Colony Management, Scientific Protocol Implementation, and the Pathology Services staff at the Wisconsin National Primate Research Center (WNPRC) for their contributions to this study. We thank BEI Resources and Brandy Russell for providing virus isolates. This work was supported by 5 R01AI132563-04 from the National Institute of Allergy and Infectious Disease and 5 P51OD011106-59 from the Office of the Director, NIH to the Wisconsin National Primate Research Center. Chelsea M. Crooks was supported by T32 AI007414 from the National Institute of Allergy and Infectious Disease.

## REFERENCES

1. Rasmussen, S. A., D. J. Jamieson, M. A. Honein, and L. R. Petersen. 2016. Zika Virus and Birth Defects--Reviewing the Evidence for Causality. N Engl J Med 374: 1981–1987.

2. Simonin, Y., F. Loustalot, C. Desmetz, V. Foulongne, O. Constant, C. Fournier-Wirth, F. Leon, J. P. Molès, A. Goubaud, J. M. Lemaitre, M. Maquart, I. Leparc-Goffart, L. Briant, N. Nagot, P. Van de Perre, and S. Salinas. 2016. Zika Virus Strains Potentially Display Different Infectious Profiles in Human Neural Cells. EBioMedicine 12: 161–169.

3. Sheridan, M. A., D. Yunusov, V. Balaraman, A. P. Alexenko, S. Yabe, S. Verjovski-Almeida, D. J. Schust, A. W. Franz, Y. Sadovsky, T. Ezashi, and R. M. Roberts. 2017. Vulnerability of primitive human placental trophoblast to Zika virus. Proc Natl Acad Sci U S A 114: E1587–E1596.

4. Sheridan, M. A., V. Balaraman, D. J. Schust, T. Ezashi, R. M. Roberts, and A. W. E. Franz. 2018. African and Asian strains of Zika virus differ in their ability to infect and lyse primitive human placental trophoblast. PLoS One 13: e0200086.

5. Smith, D. R., T. R. Sprague, B. S. Hollidge, S. M. Valdez, S. L. Padilla, S. A. Bellanca, J. W. Golden, S. R. Coyne, D. A. Kulesh, L. J. Miller, A. D. Haddow, J. W. Koehler, G. D. Gromowski, R. G. Jarman, M. T. P. Alera, I. K. Yoon, R. Buathong, R. G. Lowen, C. D. Kane, T. D. Minogue, S. Bavari, R. B. Tesh, S. C. Weaver, K. J. Linthicum, M. L. Pitt, and F. Nasar. 2018. African and Asian Zika Virus Isolates Display Phenotypic Differences Both In Vitro and In Vivo. Am J Trop Med Hyg 98: 432–444.

6. Simonin, Y., D. van Riel, P. Van de Perre, B. Rockx, and S. Salinas. 2017. Differential virulence between Asian and African lineages of Zika virus. PLoS Negl Trop Dis 11: e0005821.

7. Dowall, S. D., V. A. Graham, E. Rayner, L. Hunter, B. Atkinson, G. Pearson, M. Dennis, and R. Hewson, 2017. Lineage-dependent differences in the disease progression of Zika virus infection in type-I interferon receptor knockout (A129) mice. PLoS Negl Trop Dis 11: e0005704.

8. Jaeger, A. S., R. A. Murrieta, L. R. Goren, C. M. Crooks, R. V. Moriarty, A. M. Weiler, S. Rybarczyk, M. R. Semler, C. Huffman, A. Mejia, H. A. Simmons, M. Fritsch, J. E. Osorio, J. C. Eickhoff, S. L. O’Connor, G. D. Ebel, T. C. Friedrich, and M. T. Aliota, 2019. Zika viruses of African and Asian lineages cause fetal harm in a mouse model of vertical transmission. PLoS Negl Trop Dis 13: e0007343.

9. McDonald, E. M., N. K. Duggal, M. J. Delorey, J. Oksanish, J. M. Ritter, and A. C. Brault, 2019. Duration of seminal Zika viral RNA shedding in immunocompetent mice inoculated with Asian and African genotype viruses. Virology 535: 1–10.

10. Aliota, M. T., D. M. Dudley, C. M. Newman, E. L. Mohr, D. D. Gellerup, M. E. Breitbach, C. R. Buechler, M. N. Rasheed, M. S. Mohns, A. M. Weiler, G. L. Barry, K. L. Weisgrau, J. A. Eudailey, E. G. Rakasz, L. J. Vosler, J. Post, S. Capuano, T. G. Golos, S. R. Permar, J. E. Osorio, T. C. Friedrich, S. L. O’Connor, and D. H. O’Connor. 2016. Heterologous Protection against Asian Zika Virus Challenge in Rhesus Macaques. PLoS Negl Trop Dis 10: e0005168.

11. Koide, F., S. Goebel, B. Snyder, K. B. Walters, A. Gast, K. Hagelin, R. Kalkeri, and J. Rayner, 2016. Development of a Zika Virus Infection Model in Cynomolgus Macaques. Front Microbiol 7: 2028.

12. Rayner, J. O., R. Kalkeri, S. Goebel, Z. Cai, B. Green, S. Lin, B. Snyder, K. Hagelin, K. B. Walters, and F. Koide, 2018. Comparative Pathogenesis of Asian and African-Lineage Zika Virus in Indian Rhesus Macaque’s and Development of a Non-Human Primate Model Suitable for the Evaluation of New Drugs and Vaccines. Viruses 10:

13. Haddow, A. D., A. Nalca, F. D. Rossi, L. J. Miller, M. R. Wiley, U. Perez-Sautu, S. C. Washington, S. L. Norris, S. E. Wollen-Roberts, J. D. Shamblin, A. E. Kimmel, H. A. Bloomfield, S. M. Valdez, T. R. Sprague, L. M. Principe, S. A. Bellanca, S. S. Cinkovich, L. Lugo-Roman, L. H. Cazares, W. D. Pratt, G. F. Palacios, S. Bavari, M. L. Pitt, and F. Nasar, 2017. High Infection Rates for Adult Macaques after Intravaginal or Intrarectal Inoculation with Zika Virus. Emerg Infect Dis 23: 1274–1281.

14. Haddow, A. D., U. Perez-Sautu, M. R. Wiley, L. J. Miller, A. E. Kimmel, L. M. Principe, S. E. Wollen-Roberts, J. D. Shamblin, S. M. Valdez, L. H. Cazares, W. D. Pratt, F. D. Rossi, L. Lugo-Roman, S. Bavari, G. F. Palacios, A. Nalca, F. Nasar, and M. L. M. Pitt. 2020. Modeling mosquito-borne and sexual transmission of Zika virus in an enzootic host, the African green monkey. PLoS Negl Trop Dis 14: e0008107.

15. WHO. 2019. Zika Epidemiology Update.

16. Kasprzykowski, J. I., K. F. Fukutani, H. Fabio, E. R. Fukutani, L. C. Costa, B. B. Andrade, and A. T. L. Queiroz. 2020. A recursive sub-typing screening surveillance system detects the arising of the ZIKV African lineage in Brazil: Is there risk of a new epidemic. Int J Infect Dis

17. de Almeida, P. R., L. P. Ehlers, M. Demoliner, A. K. A. Eisen, V. Girardi, C. De Lorenzo, M. V. Bianchi, L. Mello, S. P. Pavarini, D. Driemeier, L. Sonne, and F. R. Spilki. 2019. Detection of a novel African-lineage-like Zika virus naturally infecting free-living neotropical primates in Southern Brazil.

18. Dudley, D. M., M. T. Aliota, E. L. Mohr, A. M. Weiler, G. Lehrer-Brey, K. L. Weisgrau, M. S. Mohns, M. E. Breitbach, M. N. Rasheed, C. M. Newman, D. D. Gellerup, L. H. Moncla, J. Post, N. Schultz-Darken, M. L. Schotzko, J. M. Hayes, J. A. Eudailey, M. A. Moody, S. R. Permar, S. L. O’Connor, E. G. Rakasz, H. A. Simmons, S. Capuano, T. G. Golos, J. E. Osorio, T. C. Friedrich, and D. H. O’Connor. 2016. A rhesus macaque model of Asian-lineage Zika virus infection. Nat Commun 7: 12204.

19. Nguyen, S. M., K. M. Antony, D. M. Dudley, S. Kohn, H. A. Simmons, B. Wolfe, M. S. Salamat, L. B. C. Teixeira, G. J. Wiepz, T. H. Thoong, M. T. Aliota, A. M. Weiler, G. L. Barry, K. L. Weisgrau, L. J. Vosler, M. S. Mohns, M. E. Breitbach, L. M. Stewart, M. N. Rasheed, C. M. Newman, M. E. Graham, O. E. Wieben, P. A. Turski, K. M. Johnson, J. Post, J. M. Hayes, N. Schultz-Darken, M. L. Schotzko, J. A. Eudailey, S. R. Permar, E. G. Rakasz, E. L. Mohr, S. Capuano, A. F. Tarantal, J. E. Osorio, S. L. O’Connor, T. C. Friedrich, D. H. O’Connor, and T. G. Golos. 2017. Highly efficient maternal-fetal Zika virus transmission in pregnant rhesus macaques. PLoS Pathog 13: e1006378.

20. Mohr, E. L., L. N. Block, C. M. Newman, L. M. Stewart, M. Koenig, M. Semler, M. E. Breitbach, L. B. C. Teixeira, X. Zeng, A. M. Weiler, G. L. Barry, T. H. Thoong, G. J. Wiepz, D. M. Dudley, H. A. Simmons, A. Mejia, T. K. Morgan, M. S. Salamat, S. Kohn, K. M. Antony, M. T. Aliota, M. S. Mohns, J. M. Hayes, N. Schultz-Darken, M. L. Schotzko, E. Peterson, S. Capuano, J. E. Osorio, S. L. O’Connor, T. C. Friedrich, D. H. O’Connor, and T. G. Golos. 2018. Ocular and uteroplacental pathology in a macaque pregnancy with congenital Zika virus infection. PLoS One 13: e0190617.

21. Hirsch, A. J., V. H. J. Roberts, P. L. Grigsby, N. Haese, M. C. Schabel, X. Wang, J. O. Lo, Z. Liu, C. D. Kroenke, J. L. Smith, M. Kelleher, R. Broeckel, C. N. Kreklywich, C. J. Parkins, M. Denton, P. Smith, V. DeFilippis, W. Messer, J. A. Nelson, J. D. Hennebold, M. Grafe, L. Colgin, A. Lewis, R. Ducore, T. Swanson, A. W. Legasse, M. K. Axthelm, R. MacAllister, A. V. Moses, T. K. Morgan, A. E. Frias, and D. N. Streblow. 2018. Zika virus infection in pregnant rhesus macaques causes placental dysfunction and immunopathology. Nat Commun 9: 263.

22. Mavigner, M., J. Raper, Z. Kovacs-Balint, S. Gumber, J. T. O’Neal, S. K. Bhaumik, X. Zhang, J. Habib, C. Mattingly, C. E. McDonald, V. Avanzato, M. W. Burke, D. M. Magnani, V. K. Bailey, D. I. Watkins, T. H. Vanderford, D. Fair, E. Earl, E. Feczko, M. Styner, S. M. Jean, J. K. Cohen, G. Silvestri, R. P. Johnson, D. H. O’Connor, J. Wrammert, M. S. Suthar, M. M. Sanchez, M. C. Alvarado, and A. Chahroudi. 2018. Postnatal Zika virus infection is associated with persistent abnormalities in brain structure, function, and behavior in infant macaques. Sci Transl Med 10:

23. Cragan, J. D., C. T. Mai, E. E. Petersen, R. F. Liberman, N. E. Forestieri, A. C. Stevens, A. Delaney, A. L. Dawson, S. R. Ellington, C. K. Shapiro-Mendoza, J. E. Dunn, C. A. Higgins, R. E. Meyer, T. Williams, K. N. Polen, K. Newsome, M. Reynolds, J. Isenburg, S. M. Gilboa, D. M. Meaney-Delman, C. A. Moore, C. A. Boyle, and M. A. Honein. 2017. Baseline Prevalence of Birth Defects Associated with Congenital Zika Virus Infection – Massachusetts, North Carolina, and Atlanta, Georgia, 2013-2014. MMWR Morb Mortal Wkly Rep 66: 219–222.

24. Reynolds, M. R., A. M. Jones, E. E. Petersen, E. H. Lee, M. E. Rice, A. Bingham, S. R. Ellington, N. Evert, S. Reagan-Steiner, T. Oduyebo, C. M. Brown, S. Martin, N. Ahmad, J. Bhatnagar, J. Macdonald, C. Gould, A. D. Fine, K. D. Polen, H. Lake-Burger, C. L. Hillard, N. Hall, M. M. Yazdy, K. Slaughter, J. N. Sommer, A. Adamski, M. Raycraft, S. Fleck-Derderian, J. Gupta, K. Newsome, M. Baez-Santiago, S. Slavinski, J. L. White, C. A. Moore, C. K. Shapiro-Mendoza, L. Petersen, C. Boyle, D. J. Jamieson, D. Meaney-Delman, M. A. Honein, and Z. P. R. C. U.S. 2017. Vital Signs: Update on Zika Virus-Associated Birth Defects and Evaluation of All U.S. Infants with Congenital Zika Virus Exposure - U.S. Zika Pregnancy Registry, 2016. MMWR Morb Mortal Wkly Rep 66: 366–373.

25. Tarantal, A. F., and A. G. Hendrickx. 1988. Prenatal growth in the cynomolgus and rhesus macaque (Macaca fascicularis and Macaca mulatta): A comparison by ultrasonography. Am J Primatol 15: 309–323.

26. Tarantal, A. F. 2005. Ultrasound Imaging in Rhesus (Macaca Mulatta) and Long-Tailed (Macaca fascicularis) Macaques. Reproductive and Research Applications. The Laboratory Primate: 317–352.

27. de Rijk, E. P. C. T., and E. Van Esch. 2008. The Macaque Placenta—A Mini-Review. Toxicologic Pathology 36: 108S–118S.

28. Moreno, G. K., C. M. Newman, M. R. Koenig, M. S. Mohns, A. M. Weiler, S. Rybarczyk, K. L. Weisgrau, L. J. Vosler, N. Pomplun, N. Schultz-Darken, E. Rakasz, D. M. Dudley, T. C. Friedrich, and D. H. O’Connor. 2020. Long-Term Protection of Rhesus Macaques from Zika Virus Reinfection. J Virol 94:

29. Mulkey, S. B., G. Vezina, D. I. Bulas, Z. Khademian, A. Blask, Y. Kousa, C. Cristante, L. Pesacreta, A. J. du Plessis, and R. L. DeBiasi. 2018. Neuroimaging Findings in Normocephalic Newborns With Intrauterine Zika Virus Exposure. Pediatr Neurol 78: 75–78.

30. Valdes, V., C. D. Zorrilla, L. Gabard-Durnam, N. Muler-Mendez, Z. I. Rahman, D. Rivera, and C. A. Nelson. 2019. Cognitive Development of Infants Exposed to the Zika Virus in Puerto Rico. JAMA Netw Open 2: e1914061.

31. Mulkey, S. B., M. Arroyave-Wessel, C. Peyton, D. I. Bulas, Y. Fourzali, J. Jiang, S. Russo, R. McCarter, M. E. Msall, A. J. du Plessis, R. L. DeBiasi, and C. Cure. 2020. Neurodevelopmental Abnormalities in Children With In Utero Zika Virus Exposure Without Congenital Zika Syndrome. JAMA Pediatr

32. Nielsen-Saines, K., P. Brasil, T. Kerin, Z. Vasconcelos, C. R. Gabaglia, L. Damasceno, M. Pone, L. M. Abreu de Carvalho, S. M. Pone, A. A. Zin, I. Tsui, T. R. S. Salles, D. C. da Cunha, R. P. Costa, J. Malacarne, A. B. Reis, R. H. Hasue, C. Y. P. Aizawa, F. F. Genovesi, C. Einspieler, P. B. Marschik, J. P. Pereira, S. L. Gaw, K. Adachi, J. D. Cherry, Z. Xu, G. Cheng, and M. E. Moreira. 2019. Delayed childhood neurodevelopment and neurosensory alterations in the second year of life in a prospective cohort of ZIKV-exposed children. Nat Med 25: 1213–1217.

33. Musso, D., A. I. Ko, and D. Baud. 2019. Zika Virus Infection - After the Pandemic. N Engl J Med 381: 1444–1457.

34. Quick, J., N. D. Grubaugh, S. T. Pullan, I. M. Claro, A. D. Smith, K. Gangavarapu, G. Oliveira, R. Robles-Sikisaka, T. F. Rogers, N. A. Beutler, D. R. Burton, L. L. Lewis-Ximenez, J. G. de Jesus, M. Giovanetti, S. C. Hill, A. Black, T. Bedford, M. W. Carroll. M. Nunes, L. C. Alcantara, E. C. Sabino, S. A. Baylis, N. R. Faria, M. Loose, J. T. Simpson, O. G. Pybus, K. G. Andersen, and N. J. Loman. 2017. Multiplex PCR method for MinION and Illumina sequencing of Zika and other virus genomes directly from clinical samples. Nat Protoc 12: 1261–1276.

35. Aliota, M. T., D. M. Dudley, C. M. Newman, J. Weger-Lucarelli, L. M. Stewart, M. R. Koenig, M. E. Breitbach, A. M. Weiler, M. R. Semler, G. L. Barry, K. R. Zarbock, A. K. Haj, R. V. Moriarty, M. S. Mohns, E. L. Mohr, V. Venturi, N. Schultz-Darken, E. Peterson, W. Newton, M. L. Schotzko, H. A. Simmons, A. Mejia, J. M. Hayes, S. Capuano, M. P. Davenport, T. C. Friedrich, G. D. Ebel, S. L. O’Connor, and D. H. O’Connor. 2018. Molecularly barcoded Zika virus libraries to probe in vivo evolutionary dynamics. PLoS Pathog 14: e1006964.

36. Hansen, S. G., M. Piatak, A. B. Ventura, C. M. Hughes, R. M. Gilbride, J. C. Ford, K. Oswald, R. Shoemaker, Y. Li, M. S. Lewis, A. N. Gilliam, G. Xu, N. Whizin, B. J. Burwitz, S. L. Planer, J. M. Turner, A. W. Legasse, M. K. Axthelm, J. A. Nelson, K. Früh, J. B. Sacha, J. D. Estes, B. F. Keele, P. T. Edlefsen, J. D. Lifson, and L. J. Picker. 2013. Immune clearance of highly pathogenic SIV infection. Nature 502: 100–104.

37. Lindsey, H. S., C. H. Calisher, and J. H. Mathews. 1976. Serum dilution neutralization test for California group virus identification and serology. J Clin Microbiol 4: 503–510.

